## Supplementary Material for "Beat Perception in Polyrhythms: Time is Structured in Binary Units"

Corresponding author:

**Table S1. Polyrhythm Complexity**

| Ratio | Number of events | Number of grid points at subdivisions level | Metrical grid points on the subdivision level without an event (%) |
|---|---|---|---|
| 2:3 | 4 | 6 | 33 |
| 2:5 | 6 | 10 | 40 |
| 3:4 | 6 | 12 | 50 |
| 3:5 | 7 | 15 | 53 |
| 4:5 | 8 | 20 | 60 |
| 5:6 | 10 | 30 | 67 |

*Table S1*. We define the complexity of a polyrhythm as the relation between the number of sound events and the number of grid points at the subdivision level (i.e., the least common denominator of the two pulse trains in a polyrhythm). The first column lists the polyrhythm ratios from simple to complex. The second column describes the number of sound events for each cycle of the polyrhythm. The third column shows the number of points in the metrical grid. This number corresponds to the least common denominator and defines the subdivision level. The fourth column shows the percentages of subdivisions on the metrical grid that are silent, i.e., without any sound event. The simple 2:3 polyrhythm has less empty points on the subdivision grid (33 %) than the more complex 5:6 polyrhythm (67 %).

**Figure S1. Data Cleaning and Circular Analyses to Categorize the Tapping Responses**

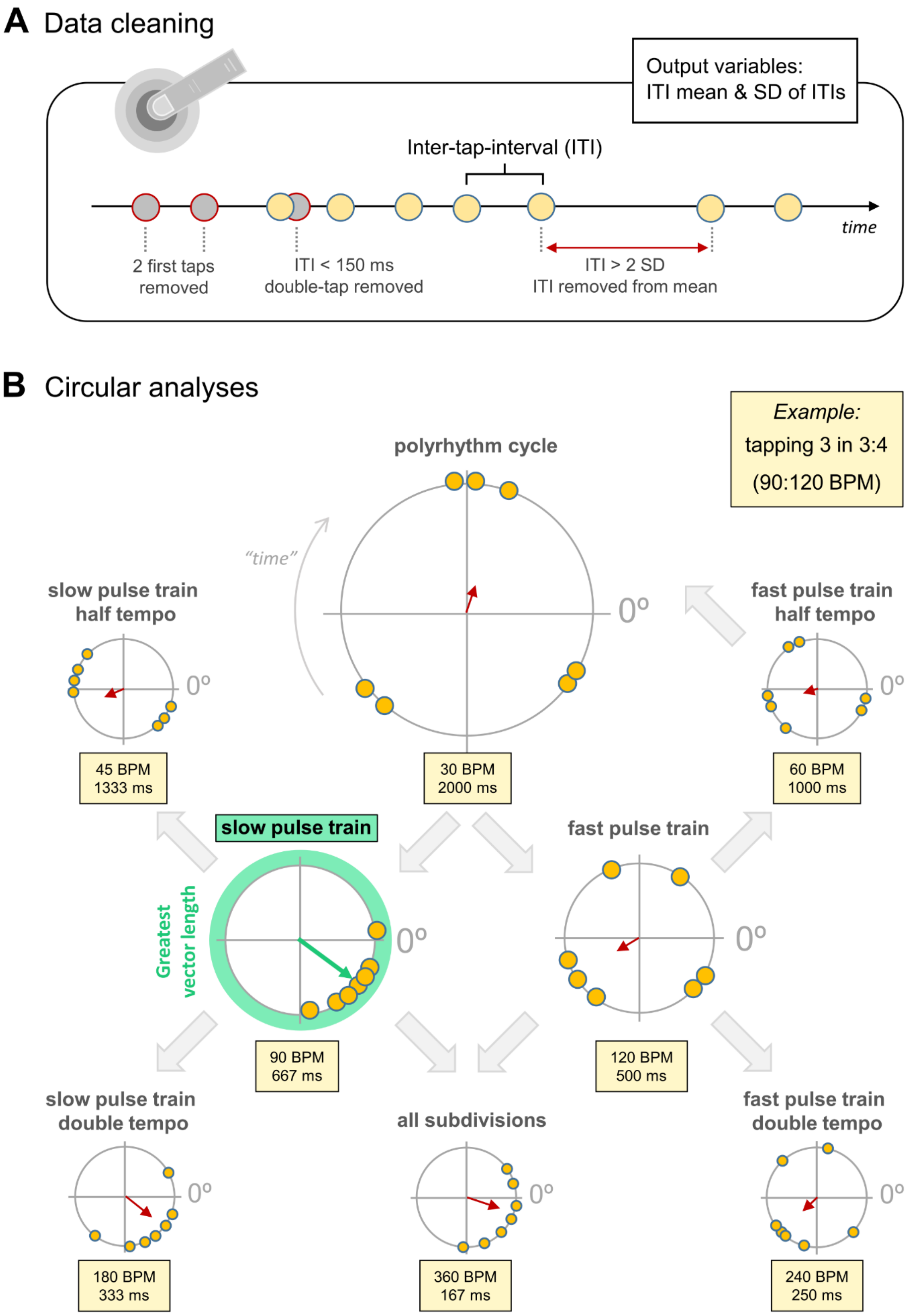

*Figure S1 A*. To obtain clean means and standard deviations of inter-tap-intervals (ITIs) in each trial, we removed the first two taps, double-taps, and ITIs beyond 2 *SD* of the mean ITI. Trials with less than five remaining taps were excluded. *Figure S1 B*. To select the metrical category that participants synchronized their taps to (example in green) and the consistency of these taps, operationalized as the vector length, the taps were transposed on a circle in relation to their "time" of occurrence within the periodicity of each metrical category. The timings of the taps were converted into angular measures (i.e., phase in radians) by calculating the modulus of each tap in reference to the tempo in milliseconds (i.e., the inter-onset interval of the full polyrhythm cycle, the slow pulse train, the double and the half tempo of the slow pulse train, the fast pulse train, the double and the half tempo of the fast pulse train, and the common subdivision level). The length of the vectors represent the consistency of the taps at each periodicity. To qualify for inclusion in a metrical category, the greatest vector length had to be not randomly distributed and significantly unimodal. This was assured by applying Rao's Spacing and Rayleigh tests. Additionally, the mean ITI had to fall within ±15 % of the metrical categories' periodicity (ms) and the SD of ITIs had to be smaller than 66 % of the metrical categories' periodicity (ms).

**Figure S2. Tempo Control Trials with 120 BPM**

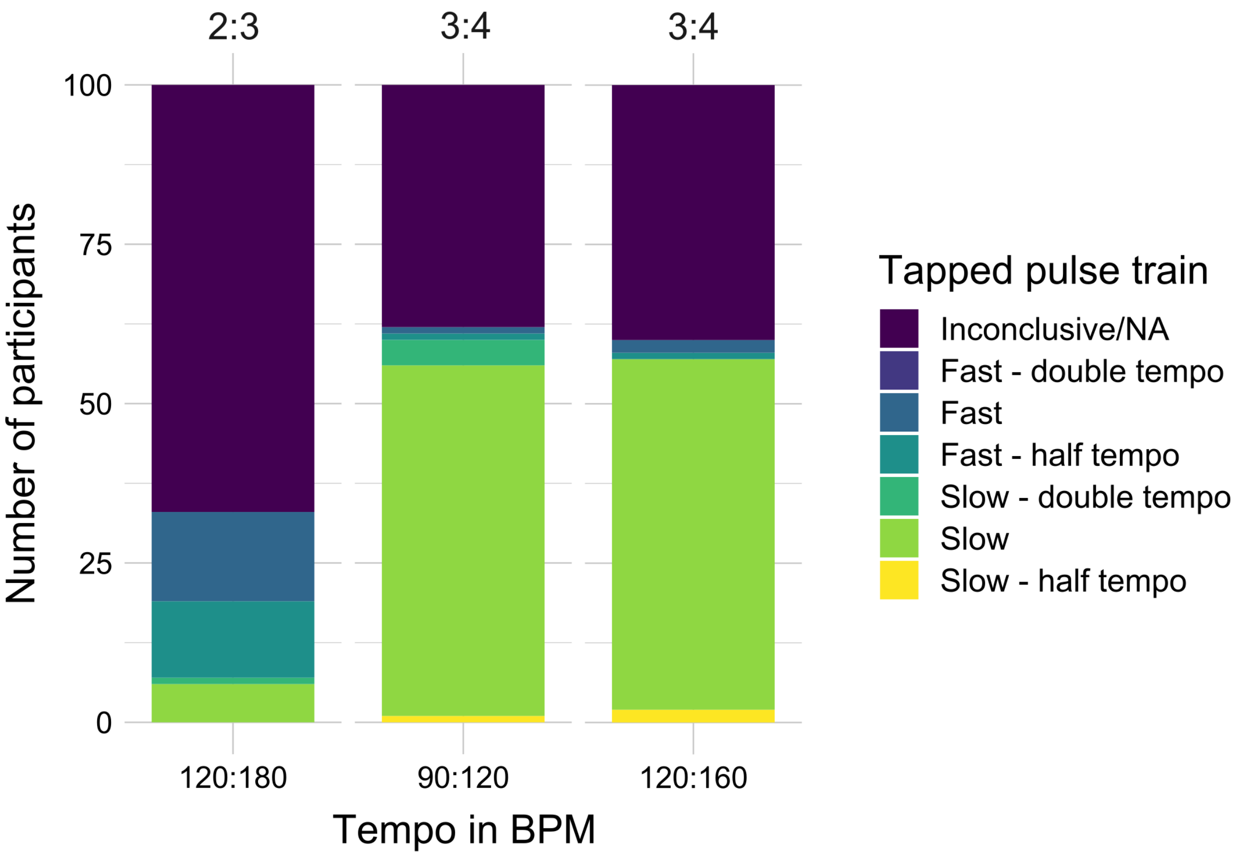


*Figure S2.* Participants' tapping responses to trials with 120 BPM in one of the pulse trains. The control reveals that responses are not driven by a tempo preference at 120 BPM. Despite the occurrence of 120 BPM in the slow pulse train of the 2:3 polyrhythm (left bar), participants tapped in time with the fast pulse train at 180 BPM or the half tempo of the fast pulse train at 90 BPM. Similarly, despite the occurrence of 120 BPM in the fast pulse train of the 3:4 polyrhythm (middle bar), participants tapped in time with the slow pulse train at 90 BPM. In both cases, participants avoided the 120 BPM pulse train in favor or the opposite pulse train which included binary subdivision grouping. Response patterns to the neighboring tempi of the 3:4 polyrhythm (middle and right bars) are almost identical suggesting that the binary subdivision grouping preference is stronger than the effect of tempo around the preferred human motor tempo of approximately 120 BPM. This is in line with the main findings of the current Tempo Experiment. Each bar in the Figure represents the number of participants tapping in time with the different metrical categories displayed in the legend on the right. The tempi of the slow and fast pulse trains are provided on the x-axis. The category "Inconclusive/NA" includes tapping to the cycle, to the subdivisions, and inconsistent tapping.

**Figure S3. Tempo Control Trials at Faster Tempi**

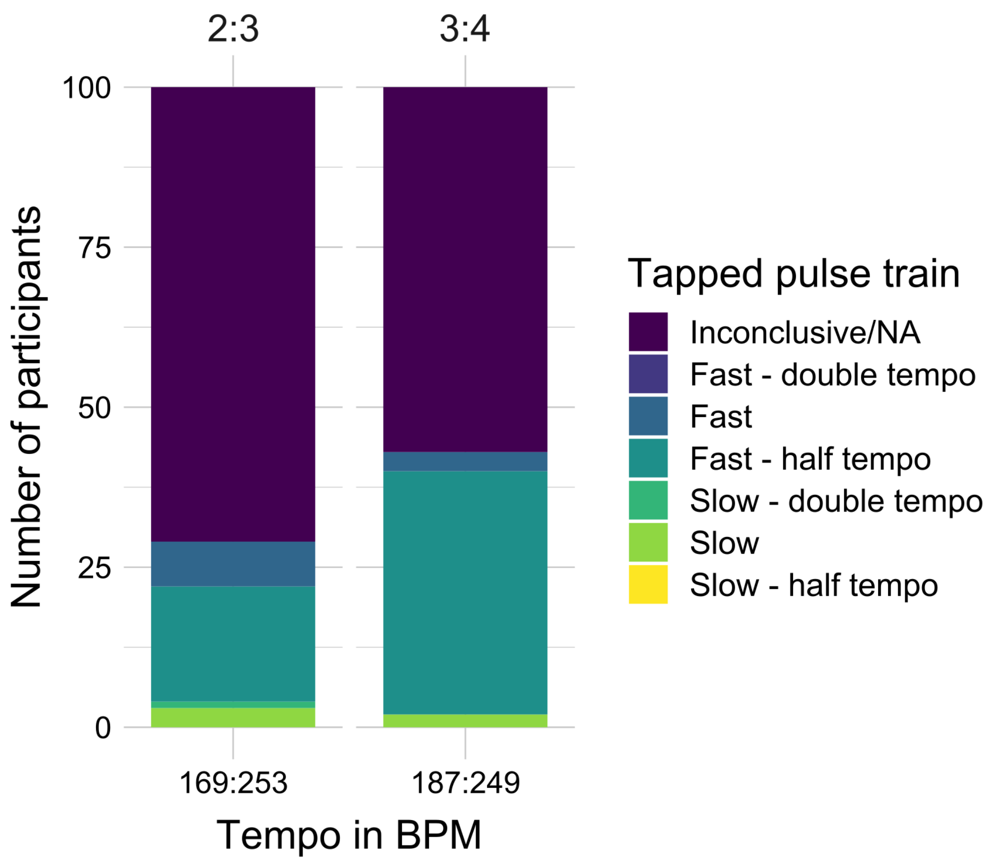


*Figure S3.* Control trials with the fast pulse train at around 250 BPM were included to cover the faster tempo range in more detail. The tapping responses show that most participants chose to tap in time with the fast pulse train at half tempo, which contain binary subdivision grouping in both cases. The 253 and 249 BPM pulse trains induced approximately 127 and 125 BPM tapping. Consistent with the main findings of the Tempo Experiment, both controls reveal that there is a preference for binary subdivision grouping. When the tempi of the physical pulse trains are faster than our preferred motor tempo, the pulse trains take the role of subdivisions and participants skip every other element in order to maintain a metrical structure containing binary subdivisions. The category "Inconclusive/NA" includes tapping to the cycle, to the subdivisions, and inconsistent tapping.

**Figure S4. Pitch Control Trials**

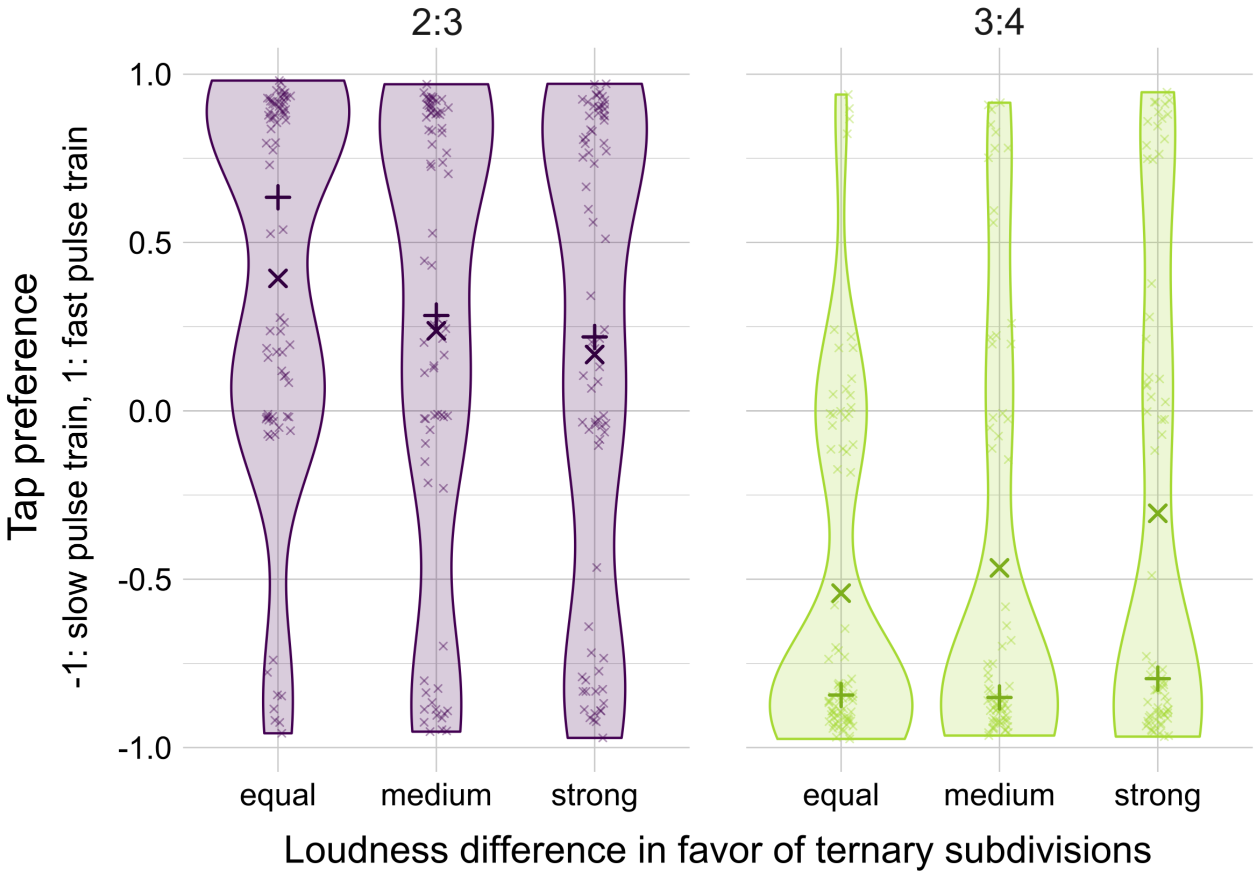


*Figure S4.* Trials controlling for the effect of amplitude in the Pitch Experiment. The pitch of both pulse trains in a polyrhythm was a C5 / 524 Hz. The amplitude manipulations followed the same method as described in the main experiment resulting in three amplitude conditions: equal loudness in both pulse trains, medium loudness difference (6 dB) in favor of the pulse train with ternary subdivisions, and strong loudness difference (12 dB) in favor of the pulse train with ternary subdivisions. The Figure depicts individual values per participant, median (+), and mean values (×) of tapping preferences with 2:3 and 3:4 polyrhythms. Values close to -1 indicate that participants consistently tapped in time with the slow pulse train, whereas values close to 1 indicate that participants consistently tapped in time with the fast pulse train. Using nonparametric pairwise comparisons (Wilcoxon signed rank test with Bonferroni corrected *p*-values), we found a difference in the individual tapping preferences between the equal and strongly modified amplitudes in both 2:3 ($Z$ = -3.46, $p$ = .002, $r$ = 0.39) and 3:4 ($Z$ = -2.58, $p$ = .030, $r$ = 0.29) polyrhythms. All other pairwise comparisons were nonsignificant ($p$-values > .07). These results indicate that there is a clear preference to tap the 3-pulse train, which is binary-subdivided (e.g., into 2 in 2:3, and into 4 in 3:4), but that this preference can be slightly shifted towards the ternary-subdivided pulse train with a strong loudness difference in favor of the ternary-subdivided pulse train. These findings complement the main Pitch Experiment by showing that the preference for synchronizing with binary-subdivided pulse trains is not only driven by pitch alternations, but already occurs in polyrhythms that use identical pitch and amplitude in the two pulse trains.
